# DOT1L suppresses nuclear RNAi originating from enhancer elements in *Caenorhabditis elegans*

**DOI:** 10.1101/320465

**Authors:** Ruben Esse, Ekaterina Gushchanskaia, Avery Lord, Alla Grishok

## Abstract

Methylation of histone H3 on lysine 79 (H3K79) by DOT1L is associated with actively transcribed genes. Earlier, we described that DOT-1.1, the *Caenorhabditis elegans* DOT1L homologue, cooperates with the chromatin-binding protein ZFP-1 (AF10 homologue) to negatively modulate transcription of highly and widely expressed target genes. Also, reduction in ZFP-1 levels has long been associated with lower efficiency of RNA interference (RNAi) triggered by exogenous double-stranded RNA (dsRNA), but the reason for this is not clear. Here, we demonstrate that DOT1L suppresses bidirectional transcription, including that producing enhancer RNAs, thereby preventing dsRNA formation and ectopic RNAi. This ectopic elevation of endogenous dsRNA may engage the Dicer complex and, therefore, limit efficiency of exogenous RNAi. Our insight provides a novel perspective on the underlying mechanisms of DOT1L function in development, neural activity, and cancer.

## INTRODUCTION

Screens for chromatin-binding factors involved in RNAi in *C. elegans* have identified the zinc finger protein ZFP-1 as a putative mediator of dsRNA-induced silencing in the nucleus (Grishok et al. 2005; Kim et al. 2005; Dudley et al. 2002). In our attempt to identify proteins interacting with ZFP-1 that could explain its role in RNAi, we previously identified the *C. elegans* homologue of the mammalian H3K79 methyltransferase DOT1L, which we named DOT-1.1 (Cecere et al. 2013). The mammalian homologue of ZFP-1, AF10, is a frequent fusion partner of MLL in chimeric proteins causing leukemia (Meyer et al. 2018), and its interaction with H3K79 methyltransferase DOT1L is critical for DOT1L recruitment to the oncogenes *HOXA9* and *MEIS1* and their subsequent activation (Okada et al. 2005).

Although DOT1L is the only H3K79 methyltransferase in mammals and H3K79 methylation is present on actively transcribed genes, inhibition of DOT1L methyltransferase activity does not result in dramatic changes in gene expression in cultured cells (Zhu et al. 2018). However, expression of specific genes, such as *HOXA9* and *MEIS1*, is strongly dependent on DOT1L, especially in leukemias induced by MLL-fusion proteins (Okada et al. 2005). The importance of H3K79 methylation in antagonizing heterochromatin formation at the *HOXA* cluster has recently been established (Chen et al. 2015), resembling pivotal studies in yeast showing that Dot1 prevents the spreading of factors associated with heterochromatin into active regions (Katan-Khaykovich and Struhl 2005; Ng et al. 2002). Importantly, knockout of DOT1L in mice results in embryonic lethality, underscoring its important developmental function (Nguyen et al. 2011a). In *Drosophila*, both DOT1L *(Grappa)* and AF10 *(Alhambra)* mutants show developmental defects (Mohan et al. 2010; Shanower et al. 2005). In *C. elegans*, we observed specific developmental abnormalities in a *zfp-1* loss-of-function mutant, such as defects in neuronal migration (Kennedy and Grishok 2014), and implicated ZFP-1 in lifespan control (Mansisidor et al. 2011). Whether the anti-silencing effect of DOT1L contributes to its developmental role is presently unknown.

We previously established that, in *C. elegans*, ZFP-1 and DOT-1.1 co-localize to promoters of highly and widely expressed genes and negatively modulate their expression during development (Cecere et al. 2013). This mechanism is likely conserved and applicable to the majority of ubiquitously expressed genes in other systems. Here, we demonstrate the role of the ZFP-1/DOT-1.1 complex in enhancer regulation. Enhancer elements are themselves transcription units, generating non-coding enhancer RNAs (eRNAs) (Li et al. 2016), and, most recently, regulation of enhancer transcription was shown to be very similar to that of protein-coding genes (Henriques et al. 2018). Indeed, we find that DOT-1.1 negatively modulates eRNA transcription, similarly to the negative modulation of active genes described earlier (Cecere et al. 2013). Moreover, we demonstrate that anti-sense transcription is also controlled by ZFP-1/DOT-1.1, and, by analyzing available genome-wide double-stranded RNA (dsRNA) and small RNA datasets, we provide evidence that regions of ZFP-1/DOT-1.1 localization, notably enhancers, are relatively depleted of anti-sense nascent transcripts, dsRNA and small RNAs.

Taken together, our observations show that elevation of endogenous dsRNA levels upon loss of ZFP-1/DOT-1.1 is most likely responsible for their apparent deficiency in responding to exogenous dsRNA (Grishok et al. 2005; Kim et al. 2005; Dudley et al. 2002). Based on our findings, we propose that negative modulation of bidirectional transcription by DOT1L does not allow dsRNA levels to reach the threshold required for inducing aberrant RNAi-based silencing and other small RNA-dependent events.

## RESULTS

### ZFP-1 and DOT-1.1 localize to predicted enhancers

Recently, the *C. elegans* genome was organized into 20 domains of different structure and activity based on chromatin modification signatures (Evans et al. 2016). We intersected coordinates of such domains with ZFP-1 and DOT-1.1 chromatin localization peaks to determine in which genomic regions the ZFP-1/DOT-1.1 complex is most dominant. More than 40% of promoter regions (domains 1 and 8 in embryos and domain 1 at L3) are enriched in ZFP-1/DOT-1.1 (Fig. 1A, Supplemental Fig. S1C), in line with our previous observations of promoter-proximal binding of the complex (Cecere et al. 2013). More interestingly, similar levels of enrichment are observed for domains corresponding to predicted intronic (domain 8 at L3) and intergenic enhancers (domain 9 in both embryos and L3 animals) (Fig. 1A,C, Supplemental Fig. S1C). These domains have chromatin signatures characteristic of enhancers, such as H3K4me1 and H3K27ac (Bonn et al. 2012). In addition, both promoters and enhancers are often devoid of nucleosomes and hence amenable to be identified by techniques assessing open chromatin regions. A recent study has identified such regions in *C. elegans* embryos and L3 animals by ATAC-seq. In this study, approximately 5000 distal noncoding regions displayed dynamic changes in chromatin accessibility between the two developmental stages. Also, several of these putative enhancers revealed enhancer activity in transgenic reporter assays (Daugherty et al. 2017). Interestingly, overlap analysis shows that both intragenic and distal ATAC-seq peaks are enriched in DOT-1.1 and ZFP-1 in embryos and L3 animals, respectively (Fig. 1B,C, Supplemental Fig. S1B). These observations raise the exciting possibility that, in addition to the previously disclosed role of ZFP-1/DOT-1.1 in the negative modulation of transcription via promoter-proximal binding, this complex may also control transcription through enhancers.

**FIGURE 1.**
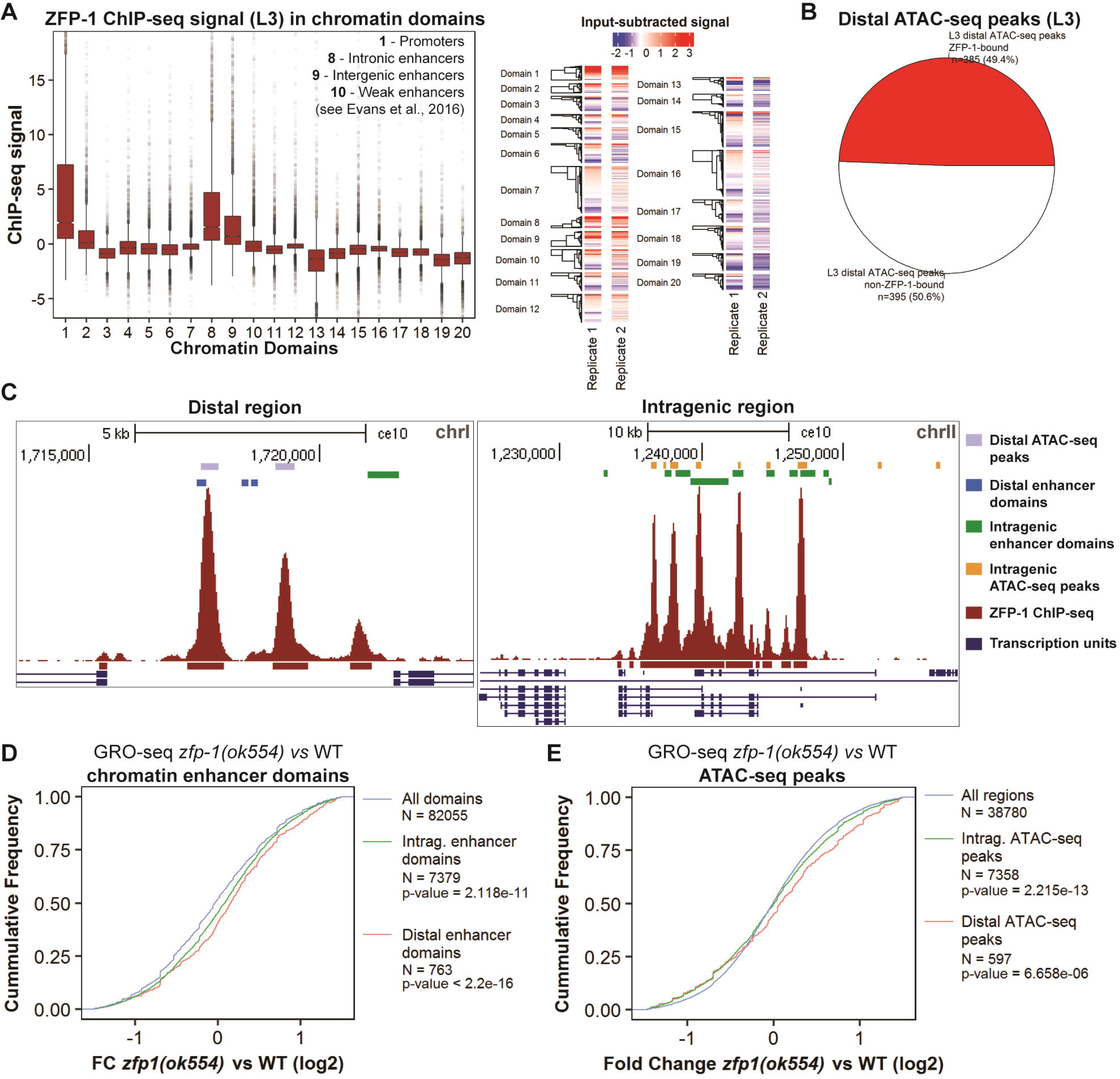
ZFP-1/DOT-1.1 negatively modulate enhancer-directed transcription. *(A)* ZFP-1 is enriched not only at chromatin domains corresponding to promoter-proximal elements, but also at predicted enhancers. Third larval stage (L3) ZFP-1 ChIP-seq coverage data in WIG format (10-bp windows) were downloaded from modENCODE (modENCODE_6213) and, for each replicate, subtracted by the corresponding input DNA (control experiment) tag density profile after normalization by sequencing depth. Chromatin domain coordinates determined by hidden Markov models were obtained from a previously published study (Evans et al. 2016) and intersected with the ZFP-1 ChIP-seq signal for each replicate. The boxplot (left panel) represents the median (medium line), first and third quartiles (box), minimum and maximum values (whiskers), 95% interval confidence of the median (notch) and outliers (dots) of the intersecting signal for each domain. Each row of the heatmap (right panel) represents a chromatin domain region and the color indicates the intersecting ZFP-1 ChIP-seq signal (averaged across replicates) as indicated in the color bar in the top. *(B)* ZFP-1 is enriched at distal enhancers identified by ATAC-seq. *C. elegans* gene coordinates (WS220/ce10) were extracted from the UCSC genome browser (http://genome.ucsc.edu/) and long (< 15 kb) gene coordinates were discarded. Gene coordinates were then intersected with open chromatin regions identified by ATAC-seq (L3) (Daugherty et al. 2017). ATAC-seq peaks overlapping with genes (flanked by 1.5 bp windows) by at least 50 bp were considered intragenic, and the remainder were considered distal. A distal ATAC-seq peak was called bound by ZFP-1 if the center base pair of at least one peak was located within its coordinates. *(C)* UCSC genome browser snapshots showing representative examples of overlap of ZFP-1 peaks with distal enhancer domains and ATAC-seq peaks (left panel) and with intragenic enhancer domains and ATAC-seq peaks (right panel). *(D)* Cumulative distribution of log2-transformed fold changes of GRO-seq RPKM (Reads Per kb per Million Reads) values at chromatin domains between *zfp-1(ok554)* and WT larvae. Raw GRO-seq reads were downloaded from NCBI GEO database (GSE47132), aligned to genome assembly WS220/ce10, normalized and counted in chromatin domains (see Materials and Methods). The p-values were determined by two-side two-sample Kolmogorov-Smirnov tests (enhancer domains compared with all chromatin domains). *(E)* Cumulative distribution of log2-transformed fold changes of GRO-seq RPKM values at genome-wide regions (all regions) and ATAC-seq peaks (Daugherty et al. 2017) between *zfp-1(ok554)* and WT larvae. Genome-wide regions were defined by dividing the genome into windows of 2.5 kb. The p-values were determined by two-side two-sample Kolmogorov-Smirnov tests (ATAC-seq peaks compared with all regions).

### ZFP-1/DOT-1.1 controls enhancer-directed transcription

In light of the observation that enhancer elements in *C. elegans* are enriched in ZFP-1/DOT-1.1, we have hypothesized that this complex could control enhancer transcription. Distal enhancers often produce eRNAs which are typically unstable and not represented in the steady-state transcript population (Li et al. 2016). This technical limitation may be circumvented by using techniques designed to assess nascent transcription, such as the global run-on sequencing (GRO-seq) method (Hah et al. 2013; Lam et al. 2013). GRO-seq data previously obtained in our lab were instrumental in providing insight into the modulatory role of ZFP-1/DOT-1.1 in transcription of genes targeted by the complex at promoter-proximal regions (Cecere et al. 2013). We re-analyzed our published data to gain a comprehensive understanding of the effect of ZFP-1/DOT1L on the regulatory genomic regions and chromatin domains revealed by recent studies (Evans et al. 2016; Daugherty et al. 2017). Nascent transcription is not limited to gene-directed transcription and is observed also at non-coding regions of the genome, including enhancers. We quantified GRO-seq coverage at genomic domains characterized by unique chromatin signatures and functional annotation (Evans et al. 2016) in wild-type (WT) and *zfp-1* mutant *(zfp-1(ok554))* L3 animals. In this mutant, ZFP-1 is C-terminally truncated and does not interact with DOT-1.1, leading to decreased abundance of DOT-1.1 at chromatin (Cecere et al. 2013). Interestingly, both intragenic and distal eRNAs were globally increased in *zfp-1* mutant animals in comparison with WT larvae (Fig. 1D, Supplemental Fig. S1D). Analysis of GRO-seq data at distal ATAC-seq peaks also revealed a global increase in transcription *zfp-1* mutant animals compared with WT larvae (Fig. 1E, Supplemental Fig. S1E). These observations indicate that, in addition to negatively modulating target gene transcription via promoter-proximal binding, the ZFP-1/DOT-1.1 complex also regulates production of RNAs derived from both intragenic and distal enhancers.

### Ectopic bidirectional transcription at ZFP-1/DOT-1.1 enhancer-containing target genes upon loss of ZFP-1

Intragenic enhancer transcription has been suggested to interfere with host gene transcription (Cinghu et al. 2017). In light of the elevation of intragenic enhancer transcription observed in the *zfp-1* mutant, we have analyzed cumulative GRO-seq coverage along gene bodies. Both sense transcription and anti-sense transcription were analyzed and each gene was considered either bound or not bound by ZFP-1, both at the promoter region and at the coding region. Overall, in the *zfp-1* mutant, genes bound by ZFP-1/DOT-1 at the promoter displayed a significant increase in both sense and anti-sense transcription, regardless of the status of enhancer signature (Supplemental Fig. S2A, top and middle panels). Transcriptional changes in the *zfp-1* mutant compared with WT at genes targeted by ZFP-1 at the coding region, but not at the promoter region, depended on the presence of enhancer chromatin signature. Target genes lacking enhancer signature displayed no overall change in transcription (Fig. 2A, top and middle panels). However, those with enhancer signature showed a significant increase in both sense (Fig. 2A, top panel) and anti-sense transcription (Fig. 2A, middle panel). These observations suggest a role for the ZFP-1/DOT-1.1 complex in preventing bidirectional transcription at target genes with intragenic enhancer signatures.

**FIGURE 2.**
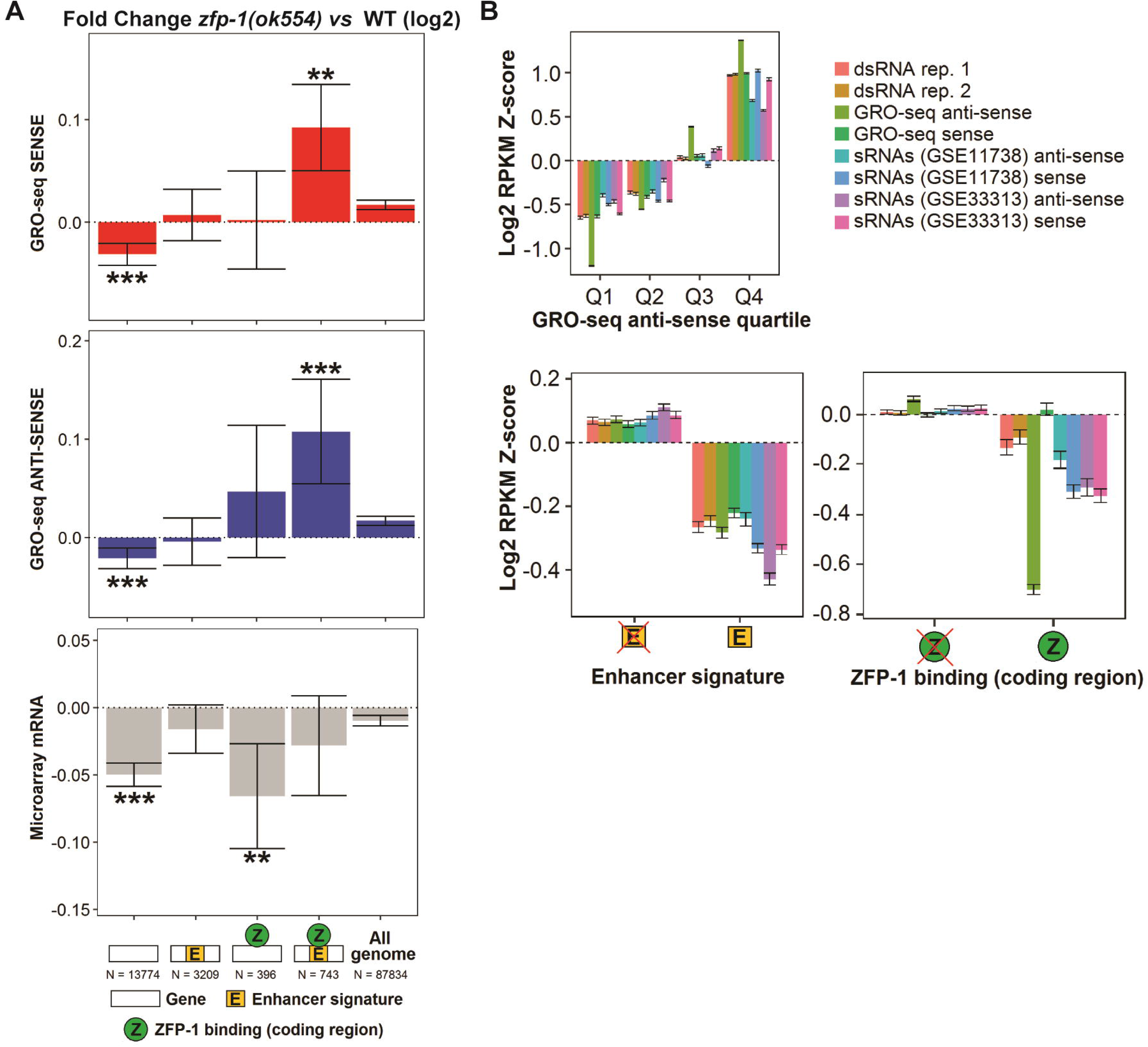
ZFP-1/DOT-1.1 prevent double-stranded RNA (dsRNA) formation at target genes with enhancer signature. *(A)* Bidirectional transcription at genes targeted by ZFP-1/DOT-1.1 at the coding region and with enhancer signature is increased in *zfp-1(ok554)* mutant animals compared with WT larvae, while the corresponding steady-state mRNA population does not overall change significantly. Genes targeted by ZFP-1/DOT-1.1 at the promoter region are not represented (see Supplemental Fig. S2). Values represent mean and 95% confidence interval. The differences between each group of genes and the group denoted as “all genome” were determined by Wilcoxon rank sum tests (** and *** denote p-values < 0.01 and < 0.001, respectively). The “all genome” group was defined by dividing the genome into windows of 2.5 kb. *(B)* The levels of nascent anti-sense RNA, small RNA and dsRNA populations originated from genes are positively correlated and are relatively depleted at enhancer-containing and ZFP-1-bound loci. Raw reads corresponding to small RNA and dsRNA populations in WT worms (L3) were downloaded from the NCBI GEO database and counted in genes (see Materials and Methods). Genes were separated in either quartiles of anti-sense GRO-seq coverage (top left panel), according to presence of enhancer signature (bottom left panel) or according to binding of ZFP-1 at the coding region (bottom right panel).

Pervasive bidirectional transcription may result in double-stranded RNA (dsRNA) formation due to complementary base pairing of sense and anti-sense transcripts (Piatek et al. 2016). We hypothesized that the increase in bidirectional transcription observed in the *zfp-1* mutant could result in formation of dsRNA. To address this hypothesis, we have resorted to publicly available small RNA (GSE11738 and GSE33313) (Batista et al. 2008; Hall et al. 2013) and dsRNA (GSE79375) (Reich et al. 2018) deep sequencing data sets obtained using L3 worms. Interestingly, small RNA and dsRNA abundances correlated positively with anti-sense GRO-seq signals (Fig. 2B, top panel, Supplemental Fig. S2B), showing that endogenous dsRNA formation stems from bi-directionally transcribed regions, ultimately leading to small RNA production. Moreover, we observed a relatively lower RNA production from enhancer-containing genes (Fig. 2B, lower left panel) and a remarkable depletion of antisense transcription at genes bound by ZFP-1 (Fig. 2B, lower right panel). These results are consistent with the proposed role of the ZFP-1/DOT-1.1 complex in inhibiting antisense transcription and the resulting dsRNA and siRNA generation.

### ZFP-1/DOT-1.1 control enhancer-containing neuronal genes

Interestingly, enhancer-containing genes targeted by ZFP-1 at the coding region did not display an overall change in mRNA levels between *zfp-1* mutant worms and WT animals (Fig. 2A, bottom panel). We hypothesized that these types of genes may exhibit tissue-specific or temporally regulated expression. In contrast to widely and highly expressed genes, developmentally regulated genes are often switched on and off in time and space, and enhancer elements play an important role in their control, including enhancers found within gene bodies. We performed pathway overrepresentation analysis of *C. elegans* genes characterized by the presence of enhancer chromatin signature and found them to be enriched in neuronal gene categories (Fig. 3A). Analysis of GRO-seq data has showed that transcription control of these genes is mediated by ZFP-1/DOT-1.1 (Fig. 2A). Therefore, it is possible that ZFP-1/DOT-1.1 control the tissue and time-specific patterning of expression of enhancer-containing neuronal genes. This observation is in line with our previous findings connecting ZFP-1 function with neuronal migration (Kennedy and Grishok 2014).

**FIGURE 3.**
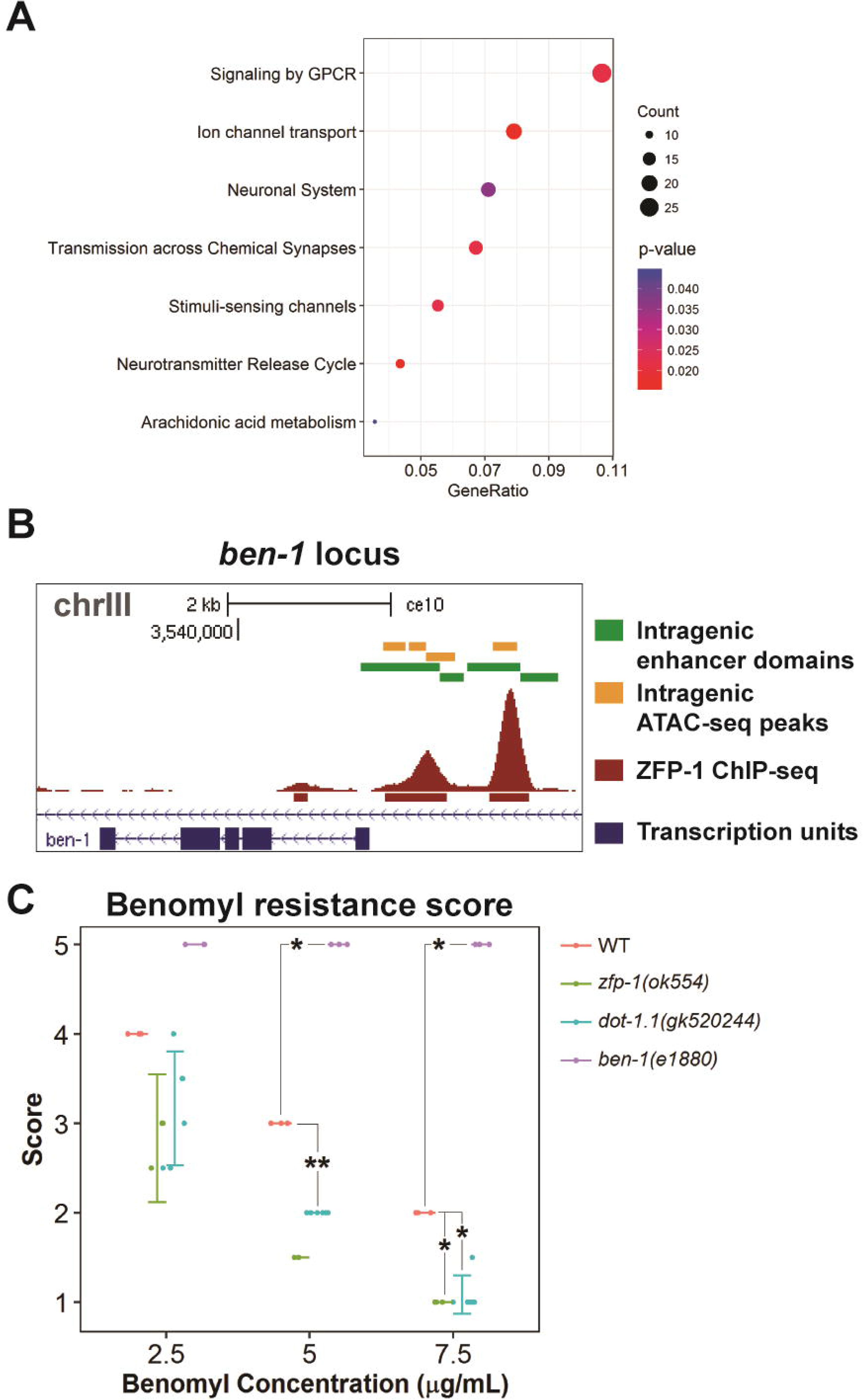
ZFP-1/DOT-1.1 control neurotransmitter receptor genes with or near enhancer elements. *(A)* Pathway overrepresentation analysis of *C. elegans* genes (WS220/ce10 assembly) with enhancer chromatin domains using the ReactomePA Bioconductor R package (Yu and He 2016). Dot sizes correspond to the number of genes in each pathway. Dot colors represent the p-values corresponding to each pathway. The x-axis shows the ratio of the number of genes in the set that are in each pathway. Some pathways include overlapping sets of genes. *(B)* UCSC genome browser snapshot showing region upstream of the *ben-1* locus characterized by the presence of enhancer chromatin signatures and ATAC-seq peaks. *ben-1* encodes a β tubulin subunit involved in chemosensory behavior and oviposition. BEN-1 is predicted to have GTP binding activity, GTPase activity, and is supposedly a structural constituent of cytoskeleton, based on protein domain information (Driscoll et al. 1989). *(C)* ZFP-1/DOT-1.1 loss-of-function mutations enhance *C. elegans* sensitivity to the antimicrotubule drug benomyl. *ben-1* loss-of-function mutations are dominant suppressors of benomyl-induced paralysis (Driscoll et al. 1989). Resistance to benomyl in the WT, *ben-1(e1880), zfp-1(ok554)* and *dot-1.1(gk520244)* alleles was scored using a 1 – 5 scale based on touch response (see Materials and Methods). Each point represents the score assigned to each tested strain in a single experiment. If displayed, error bars represent mean and 95% confidence interval. The difference between each mutant strain and the WT strain was determined by a Wilcoxon rank sum test (* and ** denote p-values < 0.05 and < 0.01, respectively).

Next, we sought out neuronal genes with well-documented mutant phenotypes as functional targets of ZFP-1/DOT-1.1. Mutations in *ben-1* are the only alleles conferring resistance to the anti-mitotic benzimidazole compound benomyl (Driscoll et al. 1989). The *ben-1* gene encodes a beta-tubulin interacting with the drug. Notably, both enhancer chromatin signature and ZFP-1/DOT-1.1 binding are present near *ben-1* (Fig. 3B). To determine whether loss-of-function in *zfp-1* or *dot-1.1* affects sensitivity to benomyl (i.e. paralysis), we subjected *zfp-1(ok554)* and *dot-1.1(gk520244)* mutants to different concentrations of the drug and found that these animals display enhanced sensitivity compared with WT worms (Fig. 3C). Therefore, it is possible that ZFP-1/DOT-1 control expression of this beta-tubulin, presumably via enhancer binding.

### Lethality of *dot-1.1* knockout is suppressed by Dicer/RDE-4/RDE-1 pathway mutations

Our attempts to generate *dot-1.1* knockout mutants in the WT background by CRISPR/Cas9 technology were not successful. Motivated by the implication of DOT1L in apoptotic cell death (Nguyen et al. 2011b; Feng et al. 2010), we have resorted to the *ced-3(n1286)* mutant strain, which is defective in the pro-apoptotic caspase CED-3 (homologue of mammalian caspase-3) (Xue et al. 1996), and have successfully generated a viable *dot-1.1(knu339)* null mutant in this mutant background (Fig. 4A). Upon outcrossing the *ced-3(n1286)* allele, we found that *dot-1.1* null animals were not viable (Fig. 4B).

**FIGURE 4.**
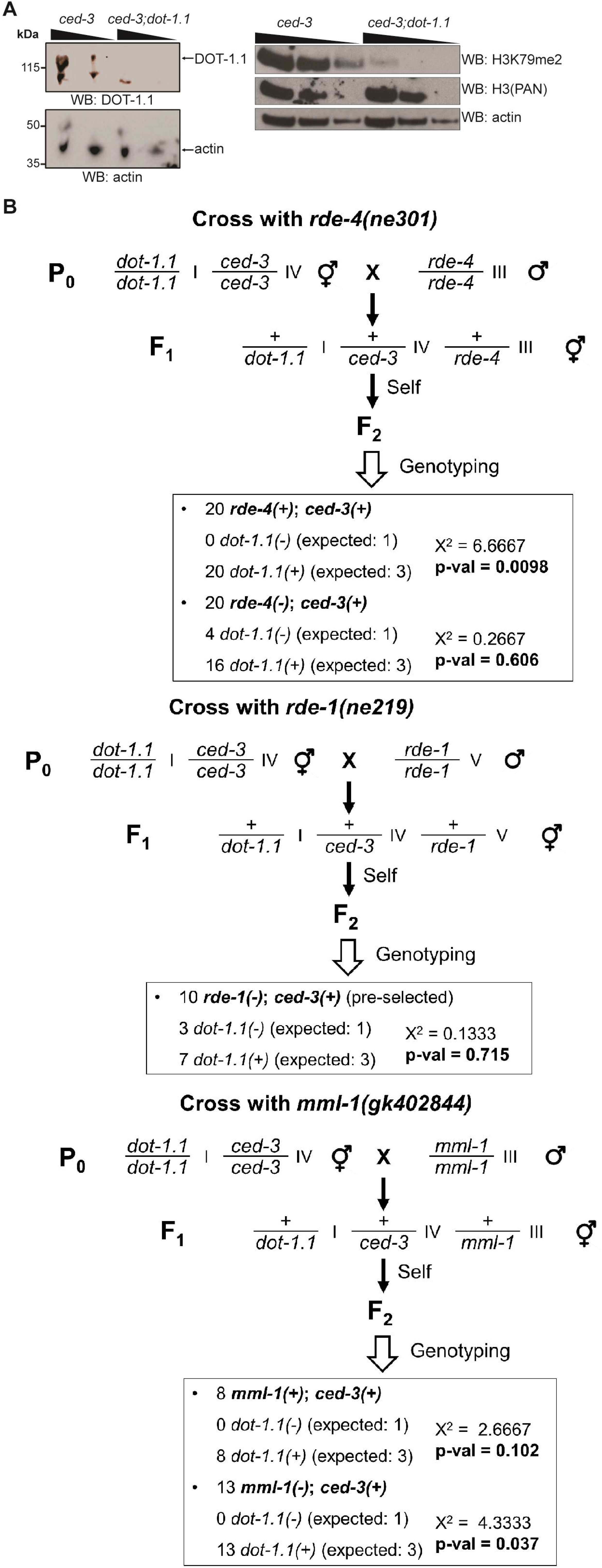
A lack of RNAi pathway components RDE-4 or RDE-1 rescues lethality of *dot-1.1* knockout. *(A)* Western blotting images showing absence of DOT-1.1 protein in a mutant strain in which the *dot-1.1* locus was deleted by CRISPR/Cas9 (left panel) and decrease in H3K79me2 levels in this mutant compared with the background *ced-3(n1286)* mutant strain (right panel). *(B)* Schematic of the genetic crosses used to determine the involvement of Dicer pathway in the lethality associated with knockout of the *dot-1.1* locus. A chi-square goodness of fit test shows significant deviation from the expected 25% for *dot-1.1(-); ced-3(+)* F2 (p-value 0.0002). Male *rde-4(ne301)* animals were crossed with *dot-1.1* mutant hermaphrodites to generate cross progeny (top panel). Hermaphrodite F1 animals were individually plated and allowed to generate self-progeny. F2 animals were genotyped for the *dot-1.1* knockout allele *(dot-1.1(-))*, and then *dot-1.1(-)* animals were genotyped for the *ced-3(n1286)* and *rde-4(ne301)* alleles *(cde-3(-)* and *rde-4(-)*, respectively). Crosses using *rde-1(ne219)* (middle panel) and *mml-1(gk402844)* (bottom panel) were similar, except that *rde-1(ne219)* homozygous animals *(rde-1(-))* were selected based on survival on *pos-1(RNAi)* feeding plates, as previously described (Tabara et al. 1999). Only F2 isolates that could be propagated indefinitely across generations were considered. Calculation of chi-square values for deviation from Mendelian ratios was performed in http://www.ihh.kvl.dk/htm/kc/popgen/genetik/applets/ki.htm. Calculation of the p-value for each chi-square test was performed in https://www.danielsoper.com/statcalc/calculator.aspx?id=11.

We then hypothesized that the *dot-1.1(knu339)* lethality phenotype could be related to the increase in dsRNA observed in *zfp-1* mutant worms. In *C. elegans* and other organisms, exogenous dsRNA is subject to processing by the Dicer complex, which cleaves it into small interfering RNAs (siRNAs) (Grishok 2005). In *C. elegans*, the first discovered RNAi-deficient mutants included *rde-1* and *rde-4* (Tabara et al. 1999). The product of the *rde-1* gene is the Argonaute protein RDE-1, which binds primary siRNAs produced by Dicer cleavage (Yigit et al. 2006). The RDE-4 protein binds long dsRNA and assists Dicer in the processing reaction (Tabara et al. 2002; Parker 2006). The RDE-4 and RDE-1 proteins are also important for the anti-viral response (Wilkins et al. 2005) and thought to have a limited role in endogenous RNAi. To determine if the RDE-4/RDE-1 pathway responds to the ectopic dsRNA accumulation seen in the *zfp-1* and *dot-1.1* mutants, we crossed the double *dot-1.1; ced-3* mutant strain with loss-of-function mutants for *rde-4* and *rde-1*. Excitingly, the double *dot-1.1(knu339); rde-4(ne301)* and *dot-1.1 (knu339); rde-1 (ne219)* mutant worms were homozygous viable and did not require the presence of the *ced-3* mutant allele for the viability (Fig. 4B, top and middle panels). In a control experiment, we crossed the double *dot-1.1; ced-3* mutant strain with a mutant unrelated to the Dicer/RDE-4/RDE-1 pathway and the lethality of *dot-1.1(knu339)* was not rescued (Fig. 4B, bottom panel). These results strongly suggest that the lethality phenotype associated with *dot-1.1* deletion is due to generation of ectopic small RNAs from excessive nuclear dsRNA.

### Ectopic endogenous dsRNA due to loss of ZFP-1/DOT-1.1 may titrate Dicer complex away from exogenous dsRNA

Genetic analyzes of RNAi in *C. elegans* initially identified genes regulating silencing in response to exogenous dsRNA (exo-dsRNA) (37). Importantly, other mutants in the endogenous RNAi ERI (Enhancer RNAi) pathway were found in screens selecting for increased response to exo-dsRNA (Simmer et al. 2002; Kennedy et al. 2004; Duchaine et al. 2006; Yigit et al. 2006). The *elt-2::GFP/lacZ* transgenic strain contains a repetitive transgenic array prone to silencing through the Dicer/RDE-4/RDE-1 pathway and also facilitated by ZFP-1 (Grishok et al. 2005). This transgene is exclusively expressed in the nuclei of intestinal cells and is silenced in ERI pathway mutants (Grishok et al. 2005), such as *eri-1(mg366)* (Fig. 5A). On the contrary, in the *dot-1.1(gk520244)* partial loss-of-function mutant background, it is readily apparent that transgenic expression is increased compared to WT (Fig. 5A). Importantly, the level of GFP expression in the double mutant *dot-1.1; eri-1* remains low, similar to that of *eri-1* alone (Fig. 5A). This result indicates that the effect of *dot-1.1* mutation (i.e. transgene de-silencing) is suppressed by the loss of ERI-1 working together with Dicer. In other words: DOT-1.1 acts upstream of ERI-1/Dicer. This result is not consistent with the longstanding belief that ZFP-1 induces RNAi in the nucleus in response to siRNAs produced by Dicer. However, it completely supports our hypothesis about the mechanism of the apparent RNAi-promoting function of ZFP-1 and DOT-1.1. In summary, our results suggest that, upon loss of ZFP-1/DOT-1.1, dsRNA produced from endogenous loci enters in the competition for the Dicer complex (Fig. 5B). Our data strongly suggest that this competition occurs in the nucleus.

**FIGURE 5.**
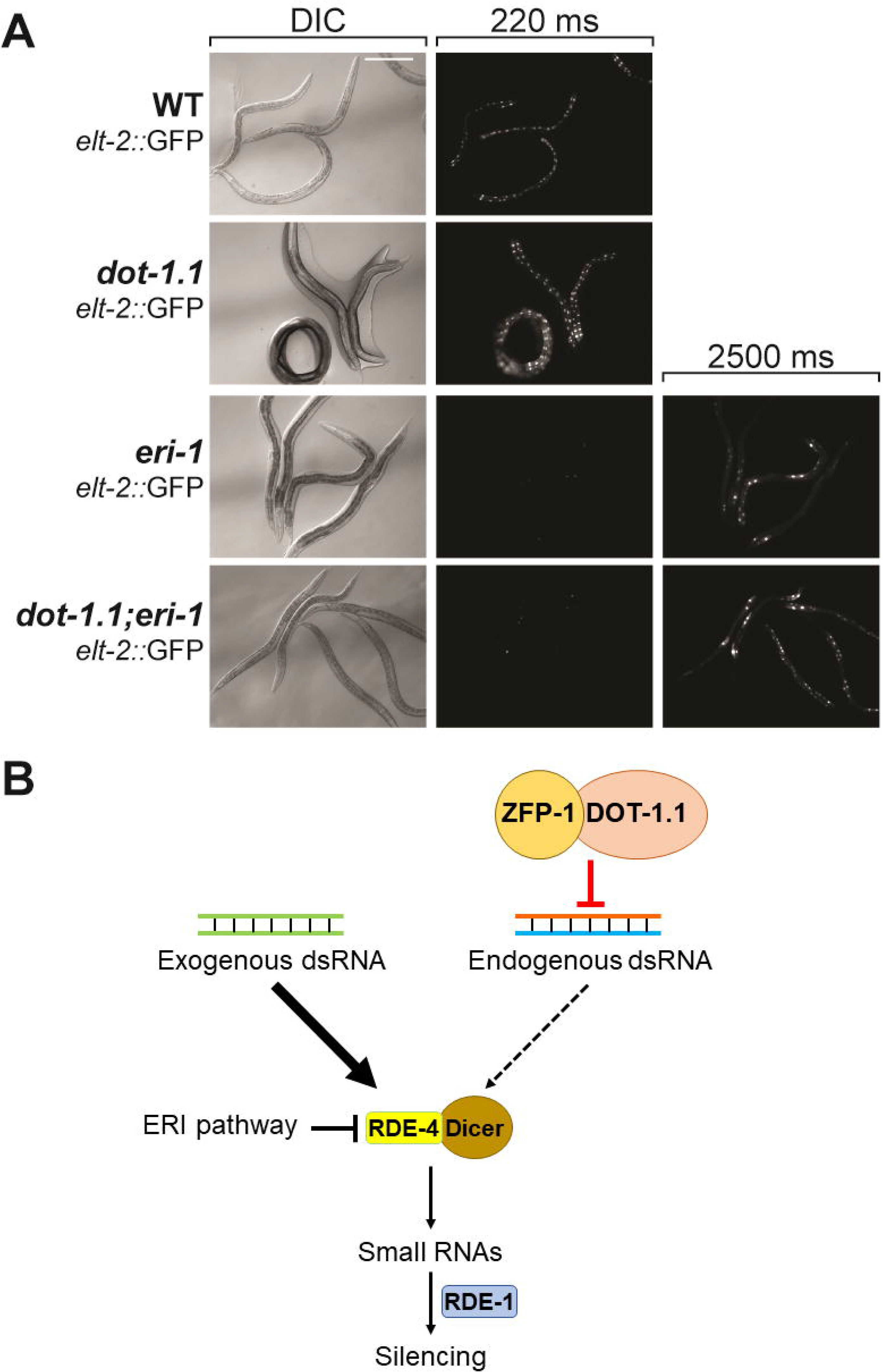
Ectopic endogenous dsRNA formed upon loss of ZFP-1/DOT-1.1 titrates the Dicer complex away from exogenous dsRNA. *(A)* In *dot-1.1(gk520244)* partial loss-of-function mutant worms, expression of the *elt-2::GFP/lacZ* transgene is increased compared with WT animals, indicating reduced efficiency of silencing by transgene-produced exogenous dsRNA. The level of GFP expression in *dot-1.1; eri-1* mutant worms is very low and similar to that observed in *eri-1* mutant worms, showing that DOT-1.1 acts upstream of ERI-1/Dicer. Scale bar (in white, upper left panel): 200 μm. More than 100 worms for each genotype were visualized, and 100% of the worms showed GFP expression presented on the images. *(B)* A model showing that ZFP-1/DOT-1.1 inhibit endogenous dsRNA production upstream of the Dicer complex, thus modulating its availability for inducing silencing by exogenous dsRNA. The ERI pathway competes for the Dicer complex downstream of ZFP-1/DOT-1.1.

## DISCUSSION

In summary, we find that, in addition to controlling transcription through promoters, as previously described (Cecere et al. 2013), the ZFP-1/DOT-1.1 complex controls enhancer-directed and antisense transcription. This finding supports the novel concept of the fluidity of enhancer/promoter states. Furthermore, we link DOT1 L-mediated transcriptional control to suppression of dsRNA formation and small RNA generation.

### Commonality between promoters and enhancers

In recent years, genomic and transcriptomic data from high-throughput sequencing studies have been shedding light on the importance of non-coding genomic regulatory regions, including enhancers. These are typically demarcated by certain histone modifications, such as H3K4me1 and H3K27ac, and open chromatin configuration. Both these features have been extensively used to predict enhancer coordinates in a variety of organisms. Importantly, new observations point to the remarkable similarity between enhancers and promoters in terms of chromatin architecture and transcriptional profile, challenging traditional binary views of enhancers and promoters (Rennie et al. 2017; Henriques et al. 2018; Mikhaylichenko et al. 2018). Now, some enhancers are thought to act as weak promoters, while, conversely, bidirectional promoters are often viewed as strong enhancers (Mikhaylichenko et al. 2018). Thus, the commonalities between enhancers and promoters may extend to how their transcriptional output is controlled by the epigenetic machinery of the cell, including chromatin writers, such as histone methyltransferases and histone acetyltransferases.

We had previously demonstrated that ZFP-1/DOT-1.1 exert a negative modulatory role on transcription of genes with promoter-proximal binding of the complex (Cecere et al. 2013). These are mostly highly and widely expressed genes not subject to spatiotemporal regulation. Here, we present evidence that the *C. elegans* DOT1L complex also modulates enhancer-directed transcription of genes subject to tissue and time-specific control of expression. Of note, the mammalian DOT1L has been implicated in processes as diverse as embryonic and postnatal hematopoiesis, proliferation of mouse embryonic stem cells, induced and natural reprogramming, cardiac development and chondrogenesis (McLean et al. 2014). Since enhancers play important roles in the regulation of these processes, it is possible that DOT1L assists promoter-enhancer cooperativity at key developmental transitions in diverse species.

### Enhancer transcription control and oncogenic function of DOT1L

Perturbation of enhancer activity is increasingly being recognized as an important player in malignant transformation (Sur and Taipale 2016). Recently, chimeric oncoproteins MLL-AF9 and MLL-AF4 were found to bind specific subsets of non-overlapping active distal enhancers in acute myeloid leukemia cell lines, and MLL-AF9-bound enhancers displayed higher levels of H3K79me2 than enhancers bound by the MLL protein alone (Prange et al. 2017). This observation can be explained by the fact that AF9, similarly to AF10, is a frequent binding partner of DOT1L and drives its localization to chromatin (Kuntimaddi et al. 2015). Therefore, it is possible that, in MLL-driven leukemias, inappropriate recruitment of DOT1L to distal regulatory elements perturbs their transcriptional output.

Homeobox loci with an indisputable role in malignant transformation, such as *HOXA9* and *MEIS1*, display both coding and non-coding transcription, including anti-sense transcripts (Popovic et al. 2008; Sessa et al. 2006), and DOT1L activity may play a yet unappreciated role in the control of such transcripts, similarly to our findings in *C. elegans*. Interestingly, a sequence conserved in vertebrate *Hox* gene introns was reported to function as enhancer element in *Drosophila* (Keegan et al. 1997; Haerry and Gehring 1996), further supporting the conservation of enhancer-control mechanisms. Moreover, the *c-myc* (Nepveu and Marcu 1986) and *N-myc* (Krystal et al. 1990) oncogenic loci display anti-sense transcription, and exciting new findings implicate DOT1L in cancers underlain by c-Myc and N-Myc activation, particularly breast cancer (Cho et al. 2015) and neuroblastoma (Wong et al. 2017). Overall, the contribution of anti-sense transcription to malignant transformation has been gaining attention in the past few years (Wenric et al. 2017; Balbin et al. 2015). Therefore, our *C. elegans* findings call for the role of anti-sense and non-coding transcription in cancer to be appreciated in the context of DOT1L activity.

### DOT1L and control of neural genes

In mammals, a set of enhancer elements characterized by the presence of the H3K4me1 mark and binding of the co-activator CBP mediate activity-dependent transcription in neurons, and this transcription is bidirectional (Kim et al. 2010; Malik et al. 2014). Also, DOT1L targets genes expressed in the cerebellum (Bovio et al. 2018) and primes neuronal layer identity in the developing cerebral cortex (Franz et al. 2018). We observed that *C. elegans* genes characterized by the presence of enhancer chromatin signatures are enriched in neuronal gene categories. Therefore, DOT1L likely operates at enhancers involved in neuronal development and activity from nematodes to humans.

### DOT1L, RNAi and cell death control

Our analysis of genome-wide GRO-seq, dsRNA, and small RNA data revealed an increase in anti-sense transcription upon loss of ZFP-1/DOT-1.1, particularly at genes with enhancer signatures, implying overlapping bidirectional transcription and formation of sense-antisense dsRNA structures. Formation of dsRNA is a common feature of multiple gene suppression phenomena known collectively as RNAi, which may be endogenous or exogenous. Importantly, production of both endogenous (endo-siRNAs) and exogenous (exo-siRNAs) small interfering RNAs requires the activity of DCR-1/Dicer. Our observations suggest that endogenous dsRNA resulting from loss of ZFP-1/DOT-1.1 competes with exogenously introduced dsRNA for DCR-1/RDE-4/RDE-1-mediated processing into small RNAs. This competition can explain the implication of ZFP-1 in RNAi (Grishok et al. 2005; Kim et al. 2005; Dudley et al. 2002).

We find that *dot-1.1* deletion mutants are not viable, which is in line with the embryonic lethality observed upon DOT1L knockout in mice (Nguyen et al. 2011a). In addition, DOT1L is required to maintain genomic and chromosomal stability, thereby preventing cell death (Giannattasio et al. 2005; Lazzaro et al. 2008; Tatum and Li 2011; Zhu et al. 2018). Interestingly, the lethality of *dot-1.1* deletion is suppressed by Dicer/RDE-4/RDE-1 pathway mutations, as well as by the mutation in the Caspase 3 gene. This suggests that small RNA generation and cell death are connected and involved in *dot-1.1* mutant lethality.

Importantly, nuclear exosome plays a key role in limiting accumulation of non-coding RNA (ncRNA), including eRNA and antisense RNA, as well as in insuring genome stability (Rothschild and Basu 2017). In this regard, ncRNA control by nuclear exosome was shown to suppress deleterious formation of RNA/DNA hybrids and dsDNA breaks at enhancers (Pefanis et al. 2015). One tantalizing possibility is that ectopic nuclear RNA species (likely siRNAs) accumulating in *dot-1.1(-)* may hybridize with enhancer DNA and expose it to nucleases, thus leading to extensive cell death and *C. elegans* lethality.

### DsRNA control by ADARs and DOT1L

It is well established that *C. elegans* ADARs (Double-Stranded RNA-Specific Adenosine Deaminases) compete for dsRNA with the DCR-1/RDE-4/RDE-1 pathway, and that *rde-4* and *rde-1* mutants suppress developmental phenotypes associated with ADAR null animals (Reich et al. 2018; Tonkin 2003; Warf et al. 2012). The difference between RNAi suppression by ADARs and that effected by ZFP-1/DOT-1.1 is that ZFP-1/DOT-1.1 prevents antisense transcription leading to dsRNA formation, whereas ADARs act after dsRNA is produced. This is consistent with the overlap between genomic regions generating dsRNA and regions of adenosine deamination (Reich et al. 2018; Reich and Bass 2018), on the one hand, and a negative correlation between ZFP-1/DOT-1.1 localization and dsRNA at coding genes loci (Fig. 2B, lower right panel), on the other. Surprisingly, in addition to negative developmental effects of ectopic dsRNA in *dot-1.1* and ADAR mutants, endogenously produced dsRNA matching highly expressed genes on chromosome arms appears to be beneficial (Reich and Bass 2018). Our analyses of GRO-seq, dsRNA and sRNA datasets (Fig. 2B, top panel) are consistent with dsRNA being largely produced from highly transcribed genomic regions in wild type worms.

Overall, our findings open numerous new directions for mechanistic research of nuclear dsRNA and RNAi in *C. elegans* and other species.

## MATERIALS AND METHODS

### Strains

Strains were maintained at 20°C under standard conditions (Brenner 1974). Bristol N2 was the WT strain used. The following strains were obtained from the Caenorhabditis Genetics Center (CGC): Bristol N2 (wild-type); VC40220, containing *dot-1.1(gk520244)* I; RB774 - *zfp-1(ok554)* III; WM49 - *rde-4(ne301)* III; VC20787, containing *mml-1(gk402844)* III; GR1373 - *eri-1(mg366)* IV; MT3002 - *ced-3(n1286)* IV; WM27 - *rde-1(ne219)* V; CB3474 - *ben-1(e1880)* III. The *rtIs11[elt-2P::GFP/lacZ]* transgenic strain was a gift from Dr. Anne Hart (Brown University). The *dot-1.1* null mutant strain COP1302, *dot-1.1 [knu339 - (pNU1092 - KO loxP::hygR::loxP)]* I; *ced-3 (n1286)* IV, in which exons one to four in the *dot-1.1* gene locus were deleted and replaced by a hygromycin insert cassette, was generated using a proprietary protocol by Knudra Transgenics. Confirmatory genotyping of the *dot-1.1(knu339)* allele was performed by PCR using the primer sets indicated in Table 1.

**Table 1.**
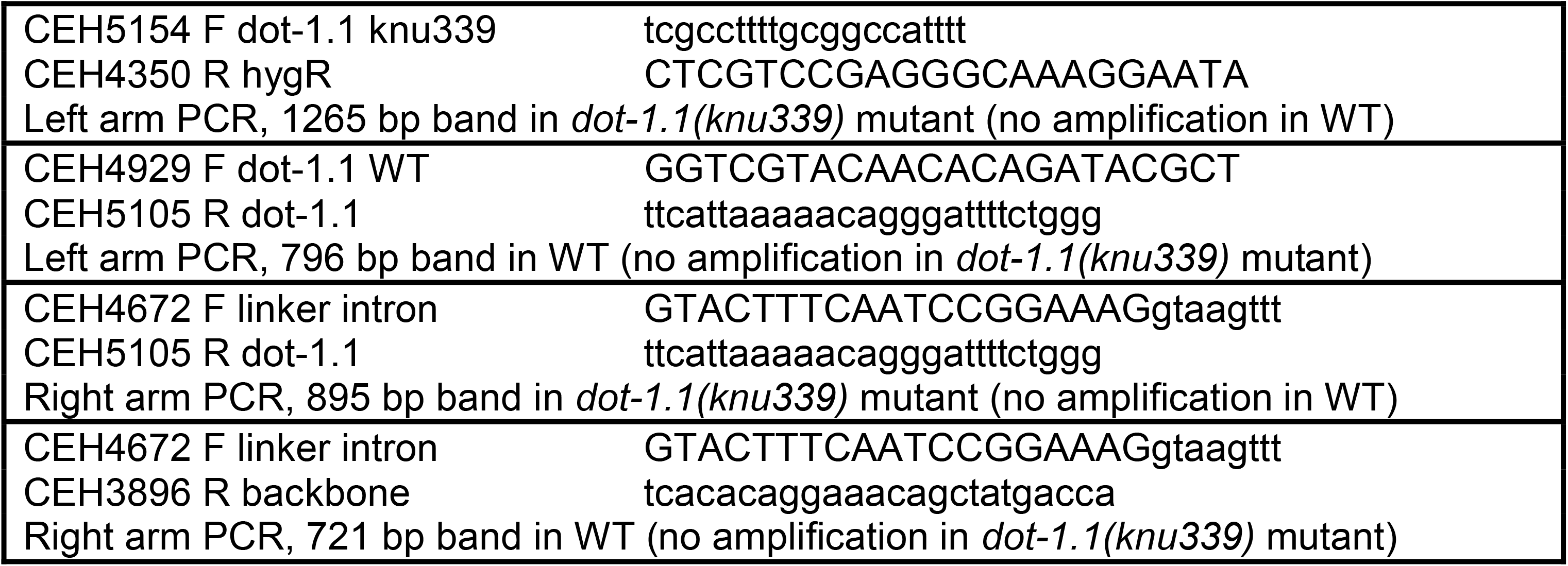
Sequences of primers used to genotype the *dot-1.1(knu339)* allele

PCR amplifications were performed with the DreamTaq™ DNA polymerase (Thermo Fisher Scientific) following the manufacturer’s instructions.

The following derivative strains were produced in this study:

AGK768: *dot-1.1(gk520244)* I, outcrossed four times from VC40220; *gk520244* (I185T) is a point mutation in the histone methyltransferase domain of DOT-1.1

AGK779: *dot-1.1(gk520244)* I; *rtIs11 (elt-2P::GFP/lacZ)* X

AGK780: *eri-1(mg366)* IV; *rtIs11 (elt-2P::GFP/lacZ)* X

AGK781: *dot-1.1(gk520244)* I; *eri-1(mg366)* IV; *rtIs11 (elt-2P::GFP/lacZ)* X

AGK782: *dot-1.1 [knu339 - (pNU1092 - KO loxP::hygR::loxP)]* I; *rde-4(ne301)* III

AGK783: *dot-1.1 [knu339 - (pNU1092 - KO loxP::hygR::loxP)]* I; *rde-1(ne219)* V

In crosses, deviations from Mendelian segregation ratios of F2 animals were calculated using chi-square goodness of fit tests (Fig. 4B).

### Western blotting

Approximately 20-25 L3 stage worms were picked directly into loading buffer (NuPAGE™ LDS Sample Buffer, Invitrogen, supplemented with NuPAGE™ Sample Reducing Agent, Invitrogen). Proteins were resolved on a precast NuPAGE Novex 4-12% Bis-Tris gel (Invitrogen) at 4 °C and transferred to a nitrocellulose membrane (0.45 μM) by semidry transfer (BioRad Trans-Blot SD transfer cell) at a constant current of 0.12 A for 1 □ hour. The blotted membrane was blocked with blocking buffer (5% non-fat dry milk in TBS-T buffer) at room temperature for 1 hour. Subsequently, it was incubated with an appropriate primary antibody overnight at 4 °C and with a secondary antibody for 2 hours at room temperature. Three washes with TBS-T buffer were made between and after the incubation with the antibodies. The membrane was developed with the SuperSignal West Femto kit (Thermo Fisher) and scanned with a KwikQuant Imager (Kindle Biosciences). The antibodies used are as follows: mouse anti-actin (Millipore, MAB1501R, 1:5000), rabbit anti-DOT-1.1 (modENCODE, SDQ3964, 1:5000), rabbit anti-H3 (Millipore, 05-928, 1:5000) and rabbit anti-H3K79me2 (Millipore, 04-835, 1:5000).

### Microscopy and immunofluorescence

The *elt-2P::GFP/LacZ* transgenic strains were grown at 16°C for at least two generations without starvation before imaging. Live animals were mounted on 2% agarose pads and examined using the Zeiss AxioImager Z1 instrument. The same exposure times were used for capturing images and comparing GFP expression in different strains.

### Benomyl sensitivity assay

Benomyl stock solution (10 mg/mL) was prepared in dimethylsulfoxide (DMSO) and stored at 4°C. Benomyl was added to melted nematode growth medium (NGM) to a concentration of 2.5 μ/mL, 5.0 μ/mL, or 7.5 μ/mL, and then NGM plates were seeded with *E. coli* OP50. Sensitivity to benomyl was scored in L3-young adult worms grown in condition plates. Scoring was done using a 1-5 scale based on touch response: 5 – complete mobility; 4 – slow response, able to back up; 3 – impaired and slow response, unable to back up; 2 – increasingly uncoordinated, very impaired; 1 – unable to effectively move away from touch. Statistical significance was determined by Wilcoxon rank sum tests.

### RNA extraction and expression profiling

Synchronized L3 larvae were washed off the plates using isotonic M9 solution. RNA was isolated in triplicate for each mutant strain by the TRIzol reagent protocol (Invitrogen) followed by miRNeasy mini column (Qiagen). RNA integrity was confirmed using a 2100 Bioanalyzer (Agilent Technologies). Samples were submitted to the BU Microarray and Sequencing Resource Core Facility for labeling and hybridization to Affymetrix GeneChip™ *C. elegans* Gene 1.0 ST arrays. Raw Affymetrix CEL files were normalized to produce gene-level expression values using the Robust Multiarray Average (RMA) (Irizarry et al. 2012) in Affymetrix Expression Console (version 1.4.1.46). The default probesets defined by Affymetrix were used to assess array quality using the Area Under the [Receiver Operating Characteristics] Curve (AUC) metric. All samples had similar quality metrics, including mean Relative Log Expression (RLE) (0.14-0.23 for all samples), and AUC values > 0.8. Fold change values were computed and log2-transformed. The results were submitted to the NCBI GEO database (GSE115677, token access slmvqiionvwfnkz).

### Analysis of chromatin domains and ATAC-seq peaks

Coordinates of chromatin domains were obtained from Evans et al., 2016 (Evans et al. 2016). DOT-1.1 ChIP-chip (mixed-stage embryo), ZFP-1 ChIP-chip (mixed-stage embryo) and ZFP-1 ChIP-seq (third larval stage, L3) peak coordinates were obtained from modENCODE (modENCODE_2970, modENCODE_3561 and modENCODE_6213 respectively). A chromatin domain region was called bound by DOT-1.1 or by ZFP-1 if the center base pair of at least one peak was located within the coordinates of the region. Putative enhancer regions at L3 were obtained by combining coordinates of domains 8, 9 and 10, annotated as intronic, intergenic and weak enhancers, respectively (Evans et al. 2016). Enhancer domains at least 1500 bp distal to any annotated transcription start site or transcription termination site were considered distal enhancer domains. Enhancer domains intersecting coordinates of genes < 15 kb by at least 50 bp were considered intragenic enhancer domains. ATAC-seq peak coordinates (Daugherty et al. 2017) were downloaded from the NCBI GEO database (GSE89608). Distal and intragenic ATAC-seq peaks were obtained as for enhancer domains. Intersections of genomic intervals were performed in R using the valr package (A. Riemondy et al. 2017). Genomic intervals obtained by the analysis were uploaded to the UCSC genome browser (https://genome.ucsc.edu/cgi-bin/hgTracks?hgS_doOtherUser=submit&hgS_otherUserName=ruben.esse&hgS_otherUserS_essionName=Esse_et_al_DOT1L_enhancers_manuscript).

### Analysis of GRO-seq data

GRO-seq data was obtained from the NCBI GEO database (GSE47132) and the reads were mapped to the *C. elegans* genome (ce10 assembly) using ChIPdig, a software application to analyze ChIP-seq data (Esse and Grishok 2017). Then, reads matching ribosomal RNA loci were removed, as described before (Cecere et al. 2013). Read counting in regions (either genes, regions or genomic bins) was performed with package GenomicAlignments (Lawrence et al. 2013); only reads with mapping quality 20 or higher were included in subsequent analyses. Regions without reads across the sample set were removed. Counts were then normalized using the TMM method, which takes RNA composition bias into account (Robinson and Oshlack 2010), using the edgeR package (Robinson et al. 2010). Coverage was expressed as RPKM (reads per kilobase per million mapped). Coverage tracks were uploaded to the UCSC genome browser (https://genome.ucsc.edu/cgi-bin/hgTracks?hgS_doOtherUser=submit&hgS_otherUserName=ruben.esse&hgS_otherUserS_essionName=Esse_et_al_DOT1L_enhancers_manuscript).

### Analyses of dsRNA and small RNA datasets

Raw reads corresponding to small RNA populations of WT animals (L3) were downloaded from the NCBI GEO database (GSE11738 and GSE33313) and aligned to genome assembly WS220/ce10. Read counting in genes and normalization was performed as for the analysis of the GRO-seq dataset. Coverage was expressed as RPKM, log2 transformed and Z-score normalized. Reads corresponding to dsRNA (GSE79375, two replicates) were processed similarly.

## ACKNOWLEDGEMENTS

Research reported in this publication was supported by the National Institute Of General Medical Sciences of the National Institutes of Health under Award Number R01GM107056. The content is solely the responsibility of the authors and does not necessarily represent the official views of the National Institutes of Health. The Caenorhabditis Genetics Center is funded by NIH Office of Research Infrastructure Programs (P40 OD010440). The BU Microarray and Sequencing Resource Core Facility is supported by the UL1TR001430 grant awarded to the BU Clinical and Translational Science Institute.

*C. elegans* strains were provided by the *C. elegans* Gene Knockout Project at the Oklahoma Medical Research Foundation, which is part of the International *C. elegans* Gene Knockout Consortium. Some strains used in this study were obtained from the Caenorhabditis Genetics Center (University of Minnesota, Minneapolis, MN). We would also like to acknowledge members of the labs of Daniel Cifuentes and Nelson Lau for discussions.

## AUTHOR CONTRIBUTIONS

R.E., E.G., A.L. and A.G. conducted experiments; R.E. analyzed genomic data; A.G. supervised the project; R.E. and A.G. wrote the manuscript.

---

**SUPPLEMENTAL FIGURE S1**. Nascent transcription at enhancers is negatively regulated by the ZFP-1/DOT-1.1 complex. *(A)* Venn diagram showing that ZFP-1 and DOT-1.1 co-localize genome-wide, as previously reported (Cecere et al. 2013). ZFP-1 and DOT-1.1 ChIP-chip peak coordinates (embryo) were downloaded from modENCODE (modENCODE_2969 and modENCODE_2970, respectively) and intersected with each other and with the full genomic space (assembly WS220/ce10). The p-value was determined by a two-sided Fisher’s exact test. *(B)* DOT-1.1 is enriched at distal enhancers identified by ATAC-seq. *C. elegans* gene coordinates (WS220/ce10) were extracted from the UCSC genome browser (http://genome.ucsc.edu/) and long (< 15 kb) gene coordinates were discarded. Gene coordinates were then intersected with open chromatin regions identified by ATAC-seq (embryo) (Daugherty et al. 2017). ATAC-seq peaks overlapping with genes (flanked by 1.5 bp windows) by at least 50 bp were considered intragenic, and the remainder were considered distal. A distal ATAC-seq peak was called bound by DOT-1.1 if the center base pair of at least one peak was located within its coordinates. *(C)* ZFP-1/DOT-1.1 complex is enriched at promoter and enhancer chromatin domains in the embryo. Chromatin domain coordinates determined by hidden Markov models and obtained from a previously published study (Evans et al. 2016) were intersected with ZFP-1 and DOT-1.1 ChIP-chip signal tracks. Each column of the heatmap (bottom panel) represents a chromatin domain region and the color indicates the intersecting ZFP-1 or DOT-1.1 ChIP-chip signal as indicated in the color bar. The boxplot (top panel) represents the median (medium line), first and third quartiles (box), minimum and maximum values (whiskers), 95% interval confidence of the median (notch) and outliers (dots) of the intersecting signal (averaged across replicates) for each domain. Boxplots showing GRO-seq RPKM (Reads Per kb per Million Reads) values at chromatin domains *(D)* and ATAC-seq peaks *(E)* for *zfp-1(ok554)* and WT third larval stage (L3) larvae. The p-values were determined by Wilcoxon rank sum tests (* and *** denote p-values < 0.05 and < 0.001, respectively). *(F)* UCSC genome browser snapshots showing representative examples of increased GRO-seq coverage in *zfp-1(ok554)* mutant animals compared with WT larvae at a distal region (left) and a genic region (right), both characterized by presence of enhancer chromatin signatures and ATAC-seq peaks.

**SUPPLEMENTAL FIGURE S2**. Ectopic double-stranded RNA (dsRNA) formation at enhancer-containing ZFP-1/DOT-1.1 targets upon loss of ZFP-1. *(A)* Global changes in sense and antisense transcription, as well as levels of steady-state transcripts, in genes bound by ZFP-1 at the promoter. Values represent mean and 95% confidence interval. The differences between each group of genes and the group denoted as “all genome” were determined by Wilcoxon rank sum tests (* and *** denote p-values < 0.05 and < 0.001, respectively). The “all genome” group was defined by dividing the genome into windows of 2.5 kb. *(B)* Sense and anti-sense transcription correlate with small RNA and dsRNA formation genome-wide. Pearson correlation coefficients are represented for each pairwise comparison, as well the p-value denoting the significance of the correlation (***, p-value < 0.001). Bivariate scatterplots with a fitted line are also represented.

