## Supplementary figures and images for "DOT1L suppresses nuclear RNAi originating from enhancer elements in *Caenorhabditis elegans*"

### Supplemental Fig. S1

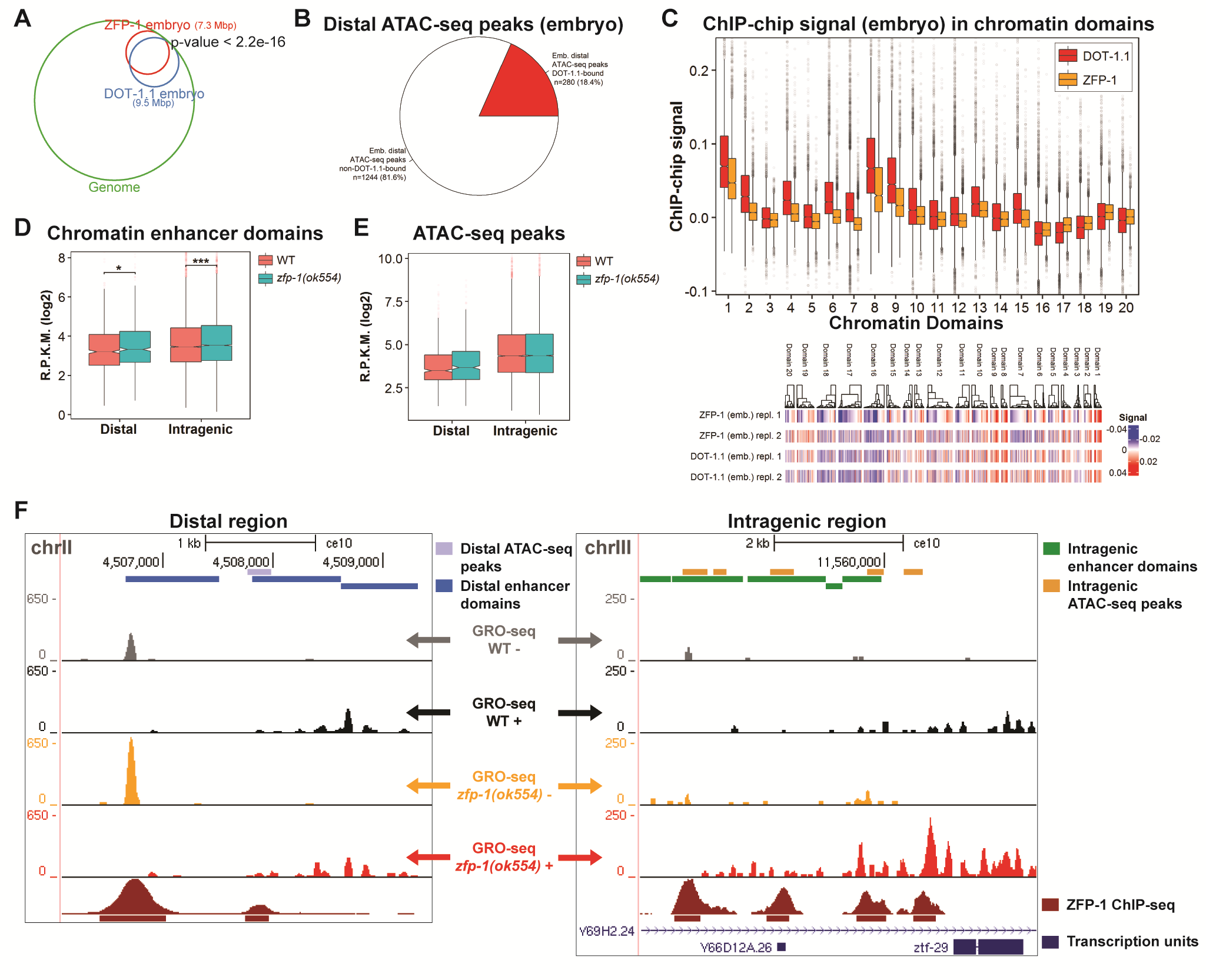

### Supplemental Fig. S2

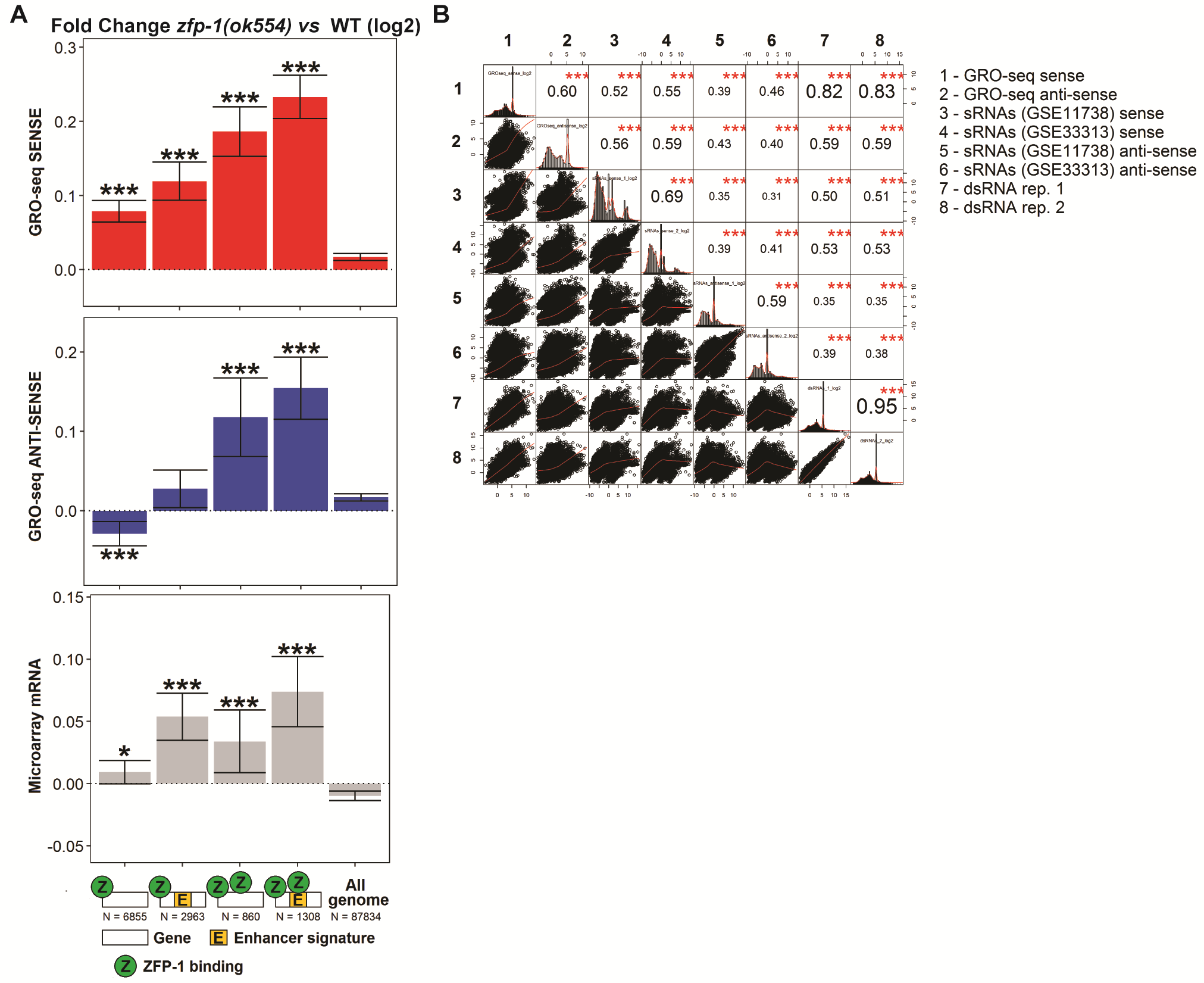
